# *In situ* structure determination using single particle cryo-electron microscopy images

**DOI:** 10.1101/2020.09.04.282509

**Authors:** Jing Cheng, Bufan Li, Long Si, Xinzheng Zhang

## Abstract

Cryo-electron microscopy (cryo-EM) tomography is a powerful tool for *in situ* structure determination. However, this method requires the acquisition of tilt series, and its time consuming throughput of acquiring tilt series severely slows determination of *in situ* structures. By treating the electron densities of non-target protein as non-Gaussian distributed noise, we developed a new target function that greatly improves the efficiency of the recognition of the target protein in a single cryo-EM image without acquiring tilt series. Moreover, we developed a sorting function that effectively eliminates the false positive detection, which not only improves the resolution during the subsequent structure refinement procedure but also allows using homolog proteins as models to recognize the target protein. Together, we developed an *in situ* single particle analysis (isSPA) method. Our isSPA method was successfully applied to solve structures of glycoproteins on the surface of a non-icosahedral virus and Rubisco inside the carboxysome. The cryo-EM data from both samples were collected within 24 hours, thus allowing fast and simple structural determination *in situ*.

## 1. Introduction

Determining the *in situ* structure of working protein machineries in their native context allows for more physiological structural information, together with identifying the interactions with other proteins nearby. One of the best technologies to determine *in situ* structures is cryo-electron tomography (Beck & Baumeister, 2016; Lučić, Leis, & Baumeister, 2008) When combined with sub-tomogram averaging technique (M. Chen et al., 2019; Leigh et al., 2019) that increases the signal noise ratio (SNR) of target protein complexes with multiple copies in the tomogram by aligning and averaging the three dimensional(3D)volume of the protein complex, *in situ* protein structures have previously been determined at sub-nanometer resolution on non-cryo-sectioning sample (Dodonova et al., 2017; Himes & Zhang, 2018; Mattei et al., 2018; Pfeffer et al., 2015; Schur, 2016; F. K. M. Schur et al., 2015; Turoňová, Schur, Wan, & Briggs, 2017; Wan et al., 2017) or nanometer resolution on cryo-sectioning sample (Bäuerlein et al., 2017; Bykov et al., 2017; Freeman Rosenzweig et al., 2017; Q. Guo et al., 2018; Mosalaganti et al., 2018). However, tomography requires the acquisition of a tilt series of the target protein complex. A tilt series typically contains more than 30 images taken at a range of tilt angles typically acquired within a time of 30 min, resulting in a slowdown of data collection throughput. The recent development of a stable sample stage in electron microscopy allows faster data collection by decreasing the waiting time of the stage (Chreifi, Chen, Metskas, Kaplan, & Jensen, 2019). However, the throughput is still dozens of times slower than the collection of single particle data.

In a single particle image, when the target protein complex is located in an *in situ* environment, the density of the target protein complex in the image is overlapped by other densities from surrounding proteins or biological molecules. The overlapping densities can be considered as noise that decreases the SNR of the image, especially at the range of low frequencies, where the shot noise is much lower than signals. The low-frequency signals exhibiting high SNR are essential for determining the initial position and orientation of the protein complex which is required by applying a conventional iterative single particle algorithm. A previous study showed that by using a high-resolution model of the target protein complex, the initial position and orientation of this protein complex can be determined from the protein background by incorporating the high-frequency signals of the target protein into the search (Rickgauer, Grigorieff, & Denk, 2017). However, the usage of the high resolution structure of the target protein renders the method less practical, since this structure is usually not available yet. A simple whitening filter was applied to both the reference and the image before calculating the correlation coefficient (Rickgauer et al., 2017), which did not take the overlapping density and SNR oscillation into account. This oscillation stems from the signal oscillation induced by the contrast transfer function (CTF). Furthermore, the shot noise in the image follows a Gaussian distribution, resulting in a smooth background in Fourier space. Therefore, CTF-like weighting has been widely used in score function (F. Guo & Jiang, 2014; Jasenko Zivanov et al., 2018). However, the overlapping densities can be regarded as part of the noise that fails to follow Gaussian distribution. How to optimize the score function when considering the noise distributed in a non-Gaussian manner remains to be investigated.

Our and other’s previous studies showed that in cases where the density of target protein complexes is overlapped by other densities, after extracting information of the initial center and orientation of the protein complex from single particle results (Zhu et al., 2018) or from sub tomogram averaging (Song et al., 2019), the structure of a protein complex can be effectively refined using traditional local refinement procedures without having to subtract overlapping densities. Both methods provide accurate initial center and orientation parameters. However, the determination of the initial center and orientation of a protein complex by incorporating a high-frequency signal with low SNR introduces false positive solutions which limits the resolution of the reconstruction. An effective method to reduce this problem and improve the resolution is urgently required.

Here, we developed a single particle-like work flow to determine the *in situ* structure of protein complexes by combining an optimized picking function to provide the initial orientation and location of a target protein and a sorting algorithm to effectively distinguish the correct solution from the false positive solutions to improve the resolution.

## 2. Results

### 2.1. Theoretical background

To localize the target protein in a crowded environment, a cross-correlation coefficient (cc) between a template of the target protein and raw cryo-EM image, had previously been applied as a picking function (Rickgauer et al., 2017). However, cryo-EM data are characterized by a frequency-dependent SNR. The template from a cryo-EM reconstruction also contains frequency-dependent noise. The SNR of the template can be calculated according to the Fourier Shell Correlation (FSC). The noise in both the cryo-EM image and the template contribute errors to the cc. Therefore, an appropriate frequency-dependent weighting function should be applied to the picking function to minimize the error in cc. In this work, the densities of the target protein in a cryo-EM image are overlapped with the densities from other surrounding proteins *in situ*. We treat the densities from surrounding proteins as noise, together with the shot noises. However, different from the shot noise, the densities from the surrounding proteins are determined by the structure factor of proteins and modified by the contrast transfer function.

Our picking function is based on cc with a frequency-dependent weight applied, which can be expressed in Fourier space as

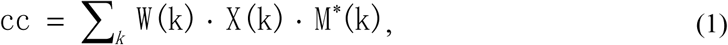

Here, *k* is the spatial frequency, *W(k)* is a weighting function, *M*\**(k)* is the conjugate complex value of the Fourier transform of a projection of a 3D template and *X(k)* is a raw image in Fourier form. Signal *S(k)* is modulated by CTF in raw image, and considering shot noise *N(k)* and protein noise which representing overlapping protein densities *PN(k), X(k)* and *M*\**(k)* are written as below

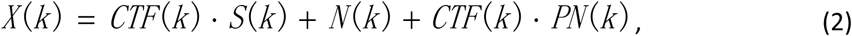

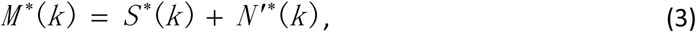

thus we can rewrite cc

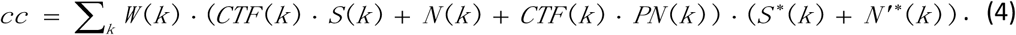

Weighting function is calculated to maximize the SNR of cc

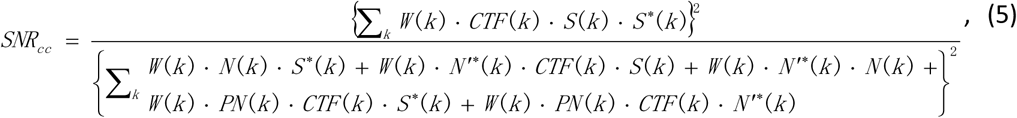

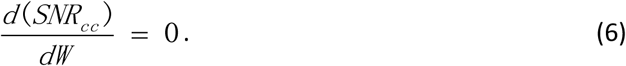

The numerator of Eq. (5) represents the square of signal of *cc*, which contributes to particle detection, and the denominator represents the variance of noise in *cc* by using the summation notation to calculate noise in cc and treating the average noise value of cc as zero since both *N* (Rosenthal & Henderson, 2003)and *PN* (Scheres, 2012) are suggested as zero-mean variations. We simplified this calculation by including in only two frequency terms (*k*_*1*_ and *k*_*2*_), and the corresponding variables are simplified to *W*_*1*_, *N*_*1*_, *N’*\*_*1*_, *PN*_*1*_, *CTF*_*1*_, and *S*_*1*_ for *W(k*_*1*_*), N(k*_*1*_*), N’(k*_*1*_*), PN(k*_*1*_*), CTF(k*_*1*_) and *S(k*_*1*_), and *W*_*2*_, *N*_*2*_, *N*^*’*\*^_*2*_, *PN*_*2*_, *CTF*_*2*_, and *S*_*2*_ for *W(k*_*2*_*), N(k*_*2*_*), N’(k*_*2*_*), PN(k*_*2*_*), CTF(k*_*2*_)

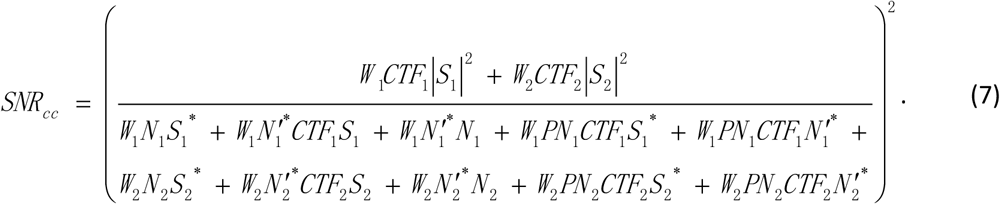

and *S(k*_*2*_) denotes the absolute value, ignoring the cross terms (noise variables are treated as random noise), and assuming that noise *N* ′ present in 3D template is much smaller than noise *N* present in 2D raw image, the result is

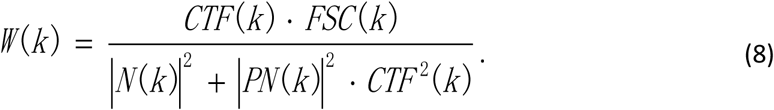

Here *FSC*(*k*) describes the Fourier Shell Correlation (FSC) between the perfect model and the 3D template, which is introduced by (Rosenthal & Henderson, 2003) as

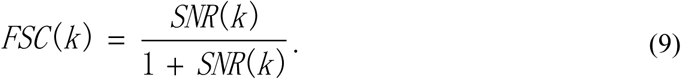

In this situation, *SNR*(*k*) equals the ratio of |*S*(*k*)|^2^ to |*N* ′(*k*)|^2^.

To remove the impacts caused by structure factor and the B-factor damping in images, phase-flipped images and projections were signal-whitened.

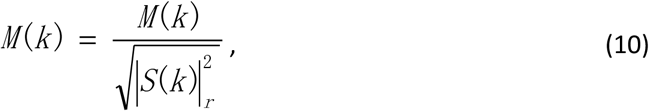

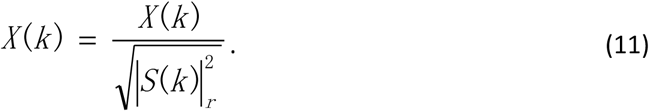

The term |*S*(*k*)|*r* ^2^ is the radially averaged signal intensity, so the weighting function applied to signal-whitened images was

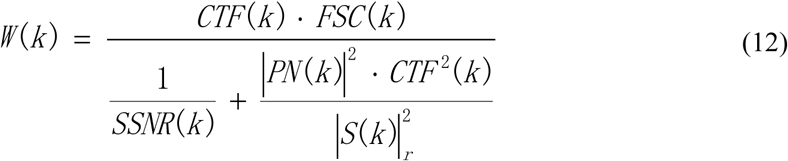

Assuming that the structural factors of different proteins are similar, the ratio of protein noise intensity to projection intensity could be approximated as a constant n, which describes how proteins are overlapped. Our optimized weighting function is simplified to

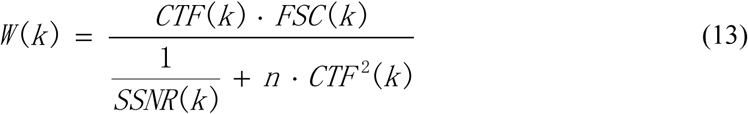

In the absence of protein noise *PN(k)*, weighting function is calculated as

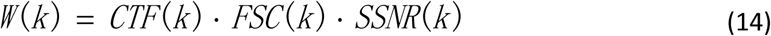

The SSNR used in the weighting function was determined from the power spectrum of the image by measures the ratio of CTF affected contrast to non-CTF affected contrast. First we fitted the non-CTF affected contrast presented in the power spectrum by fitting zero points of CTF with an exponential function. As in situ sample is usually thick (>100nm), a significant amount of none-CTF modulated noise is introduced by inelastic scattering electrons. Since inelastic electrons can be removed by energy filter which is recommended for data collection on thick sample, we did not take this into account. We calculated the differences between crest values and trough values induced by CTF oscillations on power spectrum to calculate the CTF-affected contrast. The CTF oscillation at high frequency range can be easily weakened by defocus variation in the image caused by either the distribution of proteins along the incident electron beam in thick sample or a slight unintentional tilt of the grid. Thus,, we used values below 1/8 Å^-1^ for fitting the other exponential function. The two sets of exponential function were used to calculate SSNR in the later processing.

Weighting functions from a typical cryo-EM micrograph with different values of n applied were calculated and plotted in Fig. 1. Here,SSNR was estimated from the power spectrum of the image (see Materials and Methods) and *FSC* was set to 1. An increase of n indicates more noise from overlapping protein densities, which decreases the SNR. However, the SNR at low frequency range decreases faster than that at high frequency range. Therefore, the weight of score at high frequency range increases along with an increase of n. Moreover, the shape of the peak of the oscillation of the weighting function expended in x direction with an increasing n. When n is much larger than 1/SSNR(k) (Fig. 1, n = 999) (i.e. ignoring the shot noise), our weighting function is similar to a whitening filter (Rickgauer et al., 2017). Under different conditions when n is zero (e.g. ignoring the overlapping densities), our weighting function is a dot product of *CTF*(*k*) (absolute value when applied to our signal-whitened images), *SSNR*(*k*) and *FSC*(*k*) (see Materials and Methods). Without considering the overlapping densities, similar weighting functions have been used in CisTEM (Grant, Rohou, & Grigorieff, 2018) and for CTF refinement in RELION (J. Zivanov et al., 2018). In our picking function, n is normally estimated to 3 or 4 in real cryo-EM data (see below).

**Figure 1.**
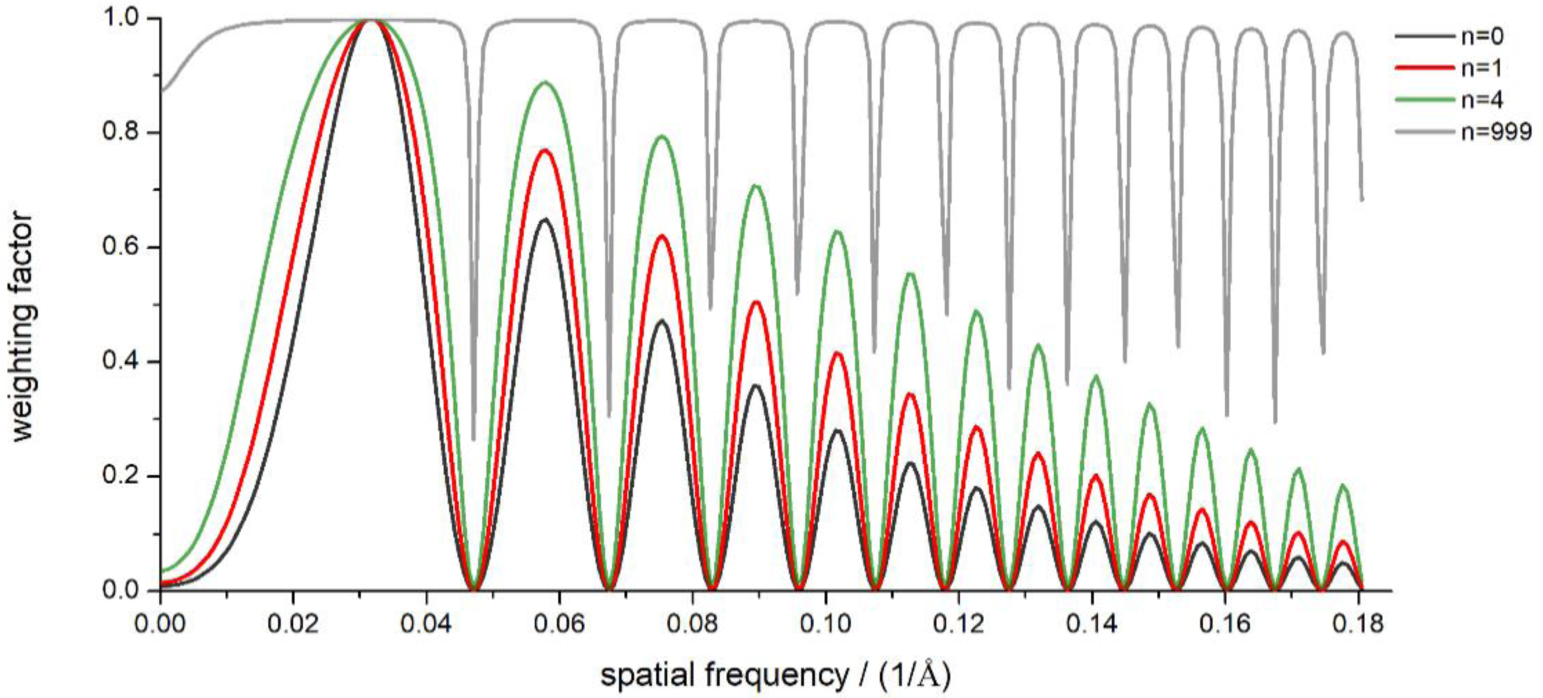
Weighting functions. Weighting factors damp and oscillate with spatial frequency (black, red, green and gray) at different n (0, 1, 4, 999).

### 2.2. Particle detection efficiency

We tested our picking function by finding different protein complexes of different sizes in cryo-EM images of different icosahedral viruses (HSV, Alphavirus, Reovirus). In these viruses, protein complexes are overlapped with densities from other proteins and genome of the virus, which mimics an *in situ* environment. High resolution single particle analysis (Yuan et al., 2018) (L. Chen et al., 2018) on these viruses ensured an accurate determination of rotational and translational parameters of each 2D image. We extracted the center and the orientation of target protein complexes on the virus and used these known parameters as positive controls. Thirty virus particle images from each of the HSV-2 and Reovirus datasets were selected for testing. Before applying a weighting function, 2D images were signal-whitened. Projections of perfect 3D model were whitened first, then the square root of each weighting function was applied to both 2D images and projections. The projections of the initial model were generated by *EMAN* (Ludtke, Baldwin, & Chiu, 1999) with the incremental rotation angle set to 5 degrees. Images and projections were binned by 2 to reduce the CPU hours. For both HSV-2 and Reovirus datasets, any results with the translational error larger than 5 pixels or the rotational error larger than 6 degrees were considered as false positive results.

In our weighting function, n regulates the ratio of protein density noise to shot noise. To pick these protein complexes, we tested different picking functions, with n ranging from 0 to 50. The results are displayed as precise-recall curves (Figs. 2B and C), where precision is the ratio of true detections to false positives and recall is the ratio of detected particles to total particles presented. When n increases from 0 to 3, the ratio of correct results increases in the datasets of HSV-2 and Reovirus. The ratio becomes steady when n is between 3 and 6. Thus, we used 3 as default value of n in the picking function. The ratio decreases when n increases further. The precise-recall curve of the whitening filter where n approaches infinity shows a much lower ratio of correct results compared with the curve of n = 3 (Figs. 2B and C).

**Figure 2.**
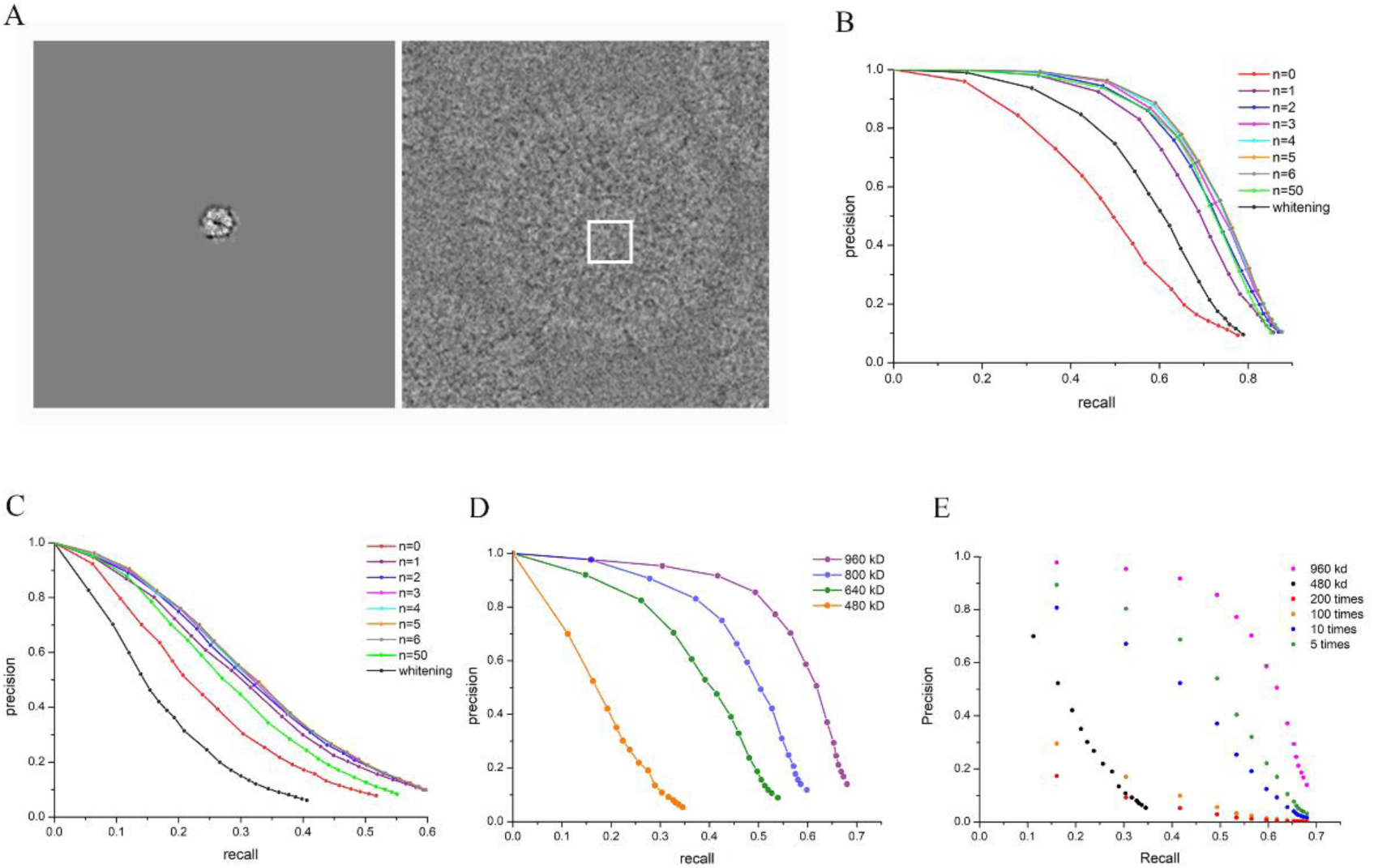
Efficiency of particle detection. (A) Left: projection of HSV-2 hexamer model. Right: HSV-2 virus particle imaged at 2.75 μm defocus with 25 electrons per Å^-2^.The location of the projection on the virus particles is indicated by a white square. (B) Precision-recall curves for detections on Reovirus datasets using model in 900 KD molecular weight at n=0 (red),1 (purple),2 (navy),3 (magenta),4 (cyan),5 (orange),6 (gray), 50 (green) and whitening filter (black). (C) Precision-recall curves for detections on HSV-2 datasets with the same processing as in (B). (D) Precision-recall curves (purple, navy, green and orange) for detections (n equals 3) on Alphaviruses using models in different molecular weights (960 kD, 800 kD, 640 kD and 480 kD). (E) Precision-recall curves for lower abundance of a 960 kD protein on Alphavirus (olive for 5 times, navy for 10 times, orange for 100 times and red for 200 times lower abundance), 960 kD at abundance of 60 copies per 90 nm * 90 nm square (magenta) and 480 kd at abundance of 60 copies per 900nm*900nm image (black).

Our picking function was tested on recognizing protein complexes of different molecular weights on Alphavirus data. As shown in Fig. 2D, the ratio of correct results versus all results decreases quickly as the size of the protein complexes decreases. This finding suggested that our picking function also generates a significant amount of incorrect particles. These false positive results are very similar to the model for picking according to the high score of our picking function, which will induce model biased feature in the reconstruction in the frequency that used for picking and noise in the structure in higher frequency in the refinement procedure.

When we are detecting a specific projection with global orientations and locations, all of the other densities (including targets at other orientations and other kinds of proteins) are contributed as noise. Therefore the SNR of low frequency signal is ruined, which makes the experiment we did on viruses different from regular single particle analysis. We think the high abundance of targets is the main superiority when compared with cellular environment and that may have an effect on detection efficiency. To investigate this, we did simulations on Alphavirus data set.

Assuming that different ratios of target protein are randomly removed from the micrographs to change the abundance of the protein, although the recall of the protein (percentage of the target proteins that are picked) remains unchanged using a fixed cutoff threshold of CC, the number of picked target protein decreases along with the decreasing of abundance. Since the background noise remains almost the same, we assume that the number of false positive result remains the same too. In such a way, we simulated the precision-recall curves of a 960kD protein with different abundances as shown in Fig. 2E. Here, when the precision drops to 0.1, the recall of the 960kD protein with 200 times decreasing of abundance is similar to that of a 480kD protein. Therefore, the size of the protein is a more important factor than the abundance.

### 2.3. Application on test data

To investigate the ability of our method to determine the protein structure in a crowded environment, we selected a part of the HSV-2 capsid as a target protein. This part is approx. 900 kD in molecular weight and consists of VP5 and VP26 trimetric capsid protein as well as the surrounding triplex (two copies of VP23 and one copy of VP19C) (Yuan et al., 2018).

We set n to 4 and used the frequencies ranging from 1/100 Å^-1^ to 1/8 Å^-1^ for particle detection. Possible locations and orientations of the target protein complex were calculated by our picking function and sorted according to the score of our picking function. After merging the results with similar orientations (within 7 degrees) in neighbor locations (within 10 pixels) into a single result, the first 500 putative target protein complexes with highest score from each virus particle were selected for further data processing, in which around 88% were false positive results according to the criteria we set in the Methods section. We performed 3D classification in RELION skipping alignment using the centers and orientations provided by our picking function. As shown in Fig. 3A, 2 out of 10 classes containing the lowest percentages of false positive result were selected, among which ∼50% were false positive results. Further classification failed to improve the ratio of correct result.

**Figure 3.**
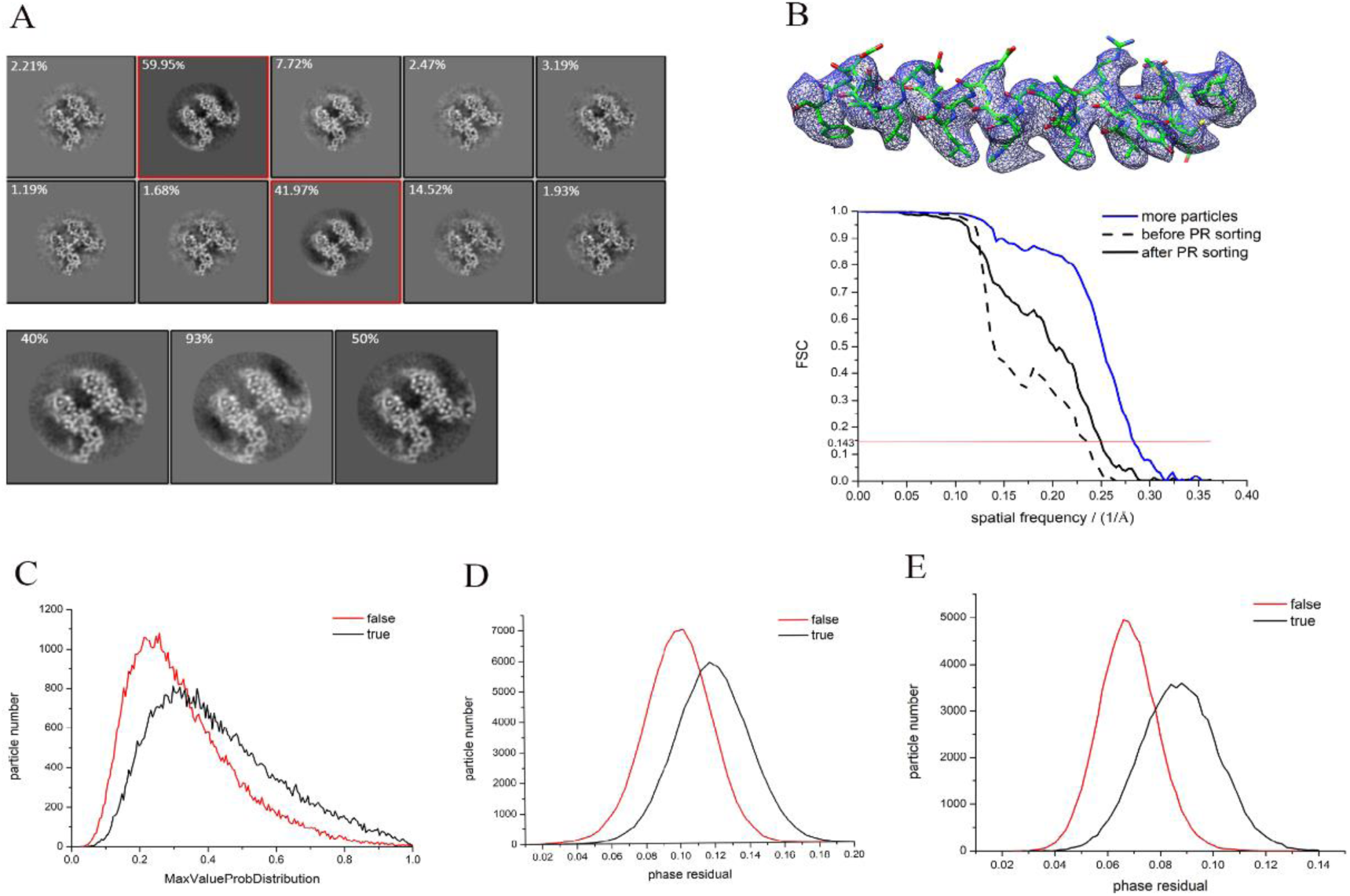
Data processing of HSV-2 hexamer. (A) Upper panel: 3D classification of raw picked positives in ten classes, the percentage of true detections in each class is shown at the top left corner. The two selected classes are indicated by squares in red. Lower panel: 3D classification of selected particles in three classes, and the percentage of true detections in each class is noted at the top left corner. (B)Upper pannel: 3D reconstruction of the HSV-2 hexamer. Lower pannel: FSC curves denote 3 reconstructions in different conditions using 2000 viral particles before PR sorting (dash) and after PR sorting (black), using 8000 virus particles and after PR sorting (navy). (C) True and false particle distribution with *MaxValueProbDistribution* term in RELION. (D) True and false particle distribution with phase residual using data from 1/30 Å^-1^ to 1/8 Å^-1^. (E) True and false particle distribution with phase residual using data from 1/8 Å^-1^ to 1/5 Å^-1^.

Next, we performed auto refinement using only local search, which resulted in a reconstruction at 4.3 Å-resolution. The FSC curve as shown in Fig.3B drops quickly at the frequency of ∼ 1/8 Å^-1^ and exhibits a shoulder around the frequency of 1/5 Å^-1^, indicating a reference bias problem (below 1/8 Å-1) and noise problem (above 1/8 Å-1) in the reconstruction. We also tried to use fewer putative target protein complexes at the top of the sorting list to increase the ratio of correct results. However, this approach also decreased the number of correct results. When we reduced the number of putative target protein complexes in the refinement procedure, the resolution of the reconstruction improved (Fig. S2) presumably due to the increase of the ratio of correct result before decreasing because of the lack of particles.

To further reduce the ratio of false positive results, we calculated the score between reference and the raw image according to refined parameters using phase residual (Methods and Materials). For this calculation, we only used the frequencies ranging from 1/8 Å^-1^ to 1/5 Å^-1^. The particles were sorted according to their score. As shown in Fig. 3E, the sorting efficiently separated the correct results from false positive results by two Gaussian-like peaks. When the sorting was based on the score from our picking function using the frequencies ranging from 1/20 Å^-1^ to 1/8 Å^-1^, the power of separation decreased markedly (Fig. 3D). This range of frequency was involved in particle picking, which performed a global search of location and orientation of the target protein on a whole virus. For instance, in a combination of translational (step size of 2.76 Å) and rotational parameters (step size of 5 degrees), 7.5 × 10^10^ possible locations of the protein complex were searched, from which the top 500 possible locations were selected by the program. Thus, each false positive result was selected from 7.5 × 10^10^ possible locations. However, further refinement using only local search improved the resolution to ∼ 4.3 Å. The local search strictly limited the possible locations. Thus, in the range of frequencies from 1/8 Å^-1^ to 1/4.3 Å^-1^, the false positive result exhibits differences from the model. Therefore, excluding the range of frequencies that involves in picking function for sorting exhibits less reference bias. In addition, the refinement only performing local search was based on maximum likelihood score function in RELION. We tested “MaxValueProbDistribution” generated in RELION as score to sort the particles; however, the correct results were barely differentiated from the false positive results (Fig. 3C). The sorting according to parameter of “NrOfSignificantSamples” exhibited similar result to that of “MaxValueProbDistribution”. Thus, it is possible that using scores different from the one used in the refinement for sort also helps to reduce the false positive detections.

After sorting by the score calculated using only the frequencies from 1/8 Å^-1^ to 1/5 Å^-1^, 40,000 particles (the top 40% particles contained ∼93% correct results) were selected. Further refinement of this dataset led to a resolution of 4.0 Å (Fig. 3B). By adding in a further 6,000 viral particles, 180,000 particles were selected after non-alignment 3D classification and sorting. The gold standard resolution was 3.7 Å. By combining with CTF refinement in RELION using our optimized weighting function, resolution was improved to 3.5 Å. The original CTF refinement procedure implanted in RELION produced a map of 3.6 Å resolution.

### 2.4. Determining protein structure using a homologous structure as a picking model

Since the expected structure from *in situ* structure determination is usually unknown, we explored the possibility of using homologous structure as a picking model. The differences between homologous structure and expected structure were treated as noise (*N*′), resulting in a different term named *FSC*_*m*_ (between the cryo-EM map of homologous structure and the expected structure) in the weighting function to replace the *FSC*.

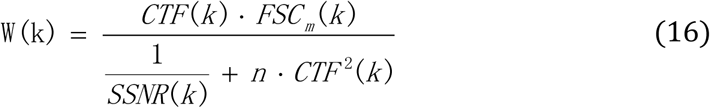

To search for a similar protein complex on HSV-2 capsid core, we used a homologous protein complex present in the HSV-1 capsid core as a model. First, we extracted the 3D model of the homologous protein complex from a 4.2 Å map of HSV-1 (Dai & Zhou, 2018). The FSC curve between protein complexes from HSV-1 and HSV-2 showed similarity in the structures with the FSC value decreasing to 0.7 at the frequency of 1/8 Å^-1^. Three million potential particles of protein complex on HSV-2 capsid were selected on the basis of the score produced by our picking function. After local 3D classification and further selection by sorting, 60,000 particles were finally selected. As shown in Fig. 4B, the local refinement resulted in a 4.0 Å resolution map. We calculated the FSC curves between this map and the corresponding 3.1 Å map from the single particle result of HSV-2 and between this map and the corresponding 4.2 Å map of HSV-1 (Dai & Zhou, 2018). As shown in Fig.4C and D, our refined structure is closer to the corresponding structure in HSV-2 than that in HSV-1. Assuming that the Å map represents a perfect map, the FSC between 3.1 Å map and our map reported a resolution of 4.0 Å using a threshold of 0.5. This result is in agreement with the resolution (4.0 Å) reported by FSC between two half maps using a threshold of 0.143. Together, these results show that the procedure we used in the *in situ* reconstruction avoids reference bias induced by homologous models.

**Figure 4.**
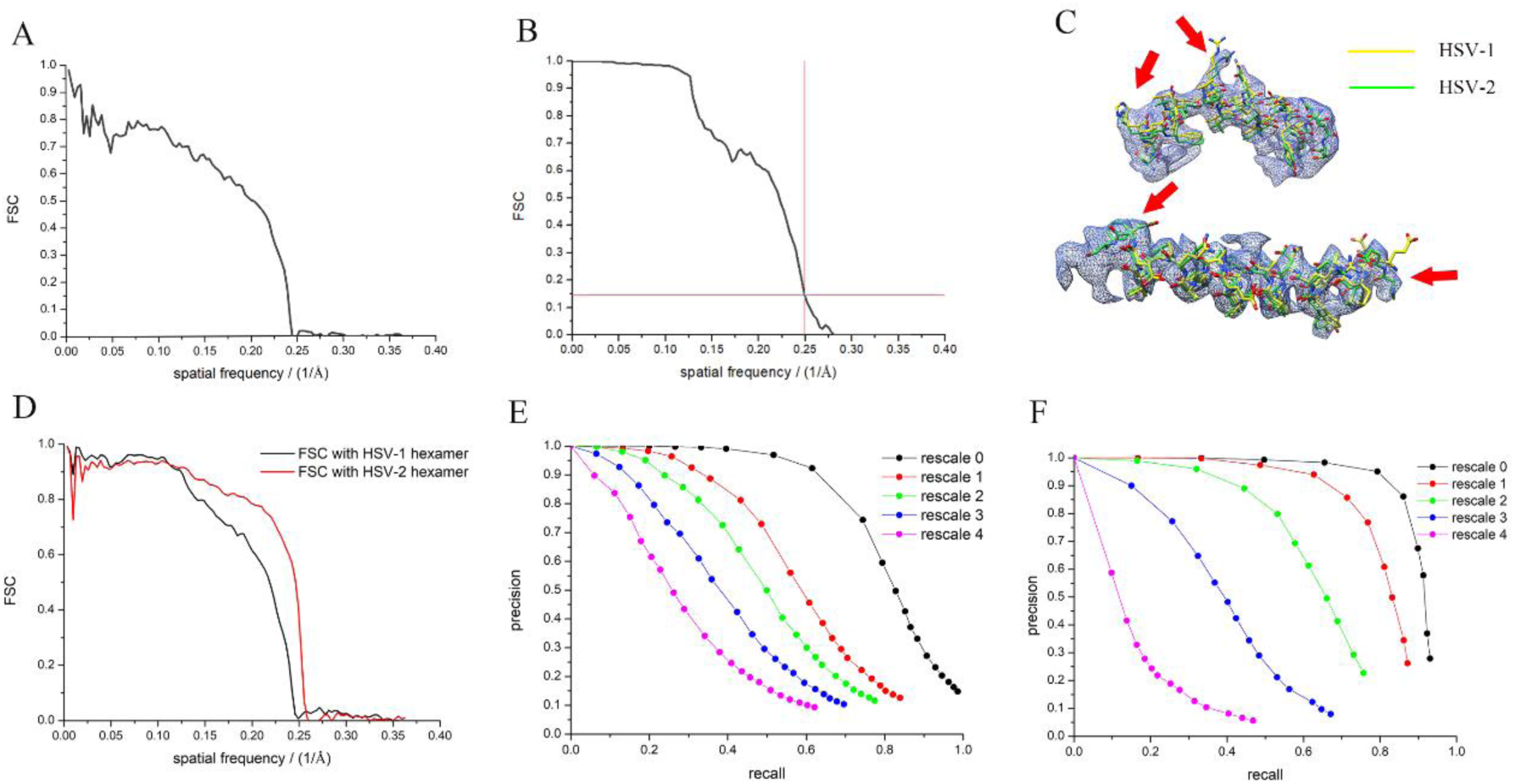
Homologous structure as picking model. (A) FSC curve of HSV-1 hexamer with HSV-2 hexamer. (B) The resolutions determined by gold standard FSC at threshold 0.143 of HSV-2 hexamer reconstructed from HSV-1 hexamer. (C) PDB of HSV-1 hexamer (yellow) and HSV-2 (green) fitting to the density of the 4.0 Å cryo-EM map, the disagreements of the two PDBs are pointed out by red arrows. (D) FSC curves show similarities between our map and HSV-1 hexamer (black), and HSV-2 hexamer (red) respectively. (E) Precision-recall curves of the ∼2 MD protein of HSV-2 at a series of scales corresponding to Fig. S3. (F) Precision-recall curves of the ∼1.8 MD protein of Reovirus at a series of scales corresponding to Fig. S3.

To evaluate the ability of finding protein complexes by homologous models of different similarity between their structures and the structure of the target protein complex, we simulated homologous models by applying different scale factors to the structure of the target protein. As shown in Fig. S3, the similarities between the re-scaled model and the original map are indicated by FSC curves. The precision-recall curves show that the ability of finding a protein complex decreases along with the reduction of the similarity between homologous model and the protein complex (Fig. 4E and F). We downloaded PDB files of different homologous proteins and calculated the potential density maps, and then calculated the FSCs between pairs of homologous maps. As shown in Fig.S4, the similarity between proteins varies greatly. These homologous proteins with high similarities can be used as picking models in our method.

### 2.5. Determining *in situ* structures at high resolution using isSPA

To evaluate our *in situ* structure determination program, we first processed a dataset of a Bunyavirus by 2D and 3D classification. As shown in Fig. 5B, only approx. 28% of the viral particles (3,140 particles) exhibited an icosahedral symmetry. Further refinement on the icosahedral viral particles resulted in an 11.8 Å map due to the flexibility. To compensate for this limitation in flexibility, one block (one pentamer and 5 surrounding hexamers) centered on a pentamer was extracted and refined (Shaikh, Hegerl, & Frank, 2003), which led to a map of 8.7 Å resolution. A centered sub-block (pentamer) and a sub-block adjacent to the pentamer that centered on a hexamer segmented from the 8.7 Å block were used as 3D models to pick the protein complex on the non-icosahedral virus particles previously excluded from data processing. Through global detection by isSPA, twenty potential solutions were selected from each virus and ∼27,000 particles in total were selected according to 3D non-alignment classification resulting in a 7.7 Å resolution map. In the search for hexamers, one hexamer from the 5 surrounding a pentamer was segmented and set as the model, and 200 potential solutions were selected from each virus. After 3D non-alignment classification, ∼87,000 particles were selected and refined to 8.3 Å resolution.

**Figure 5.**
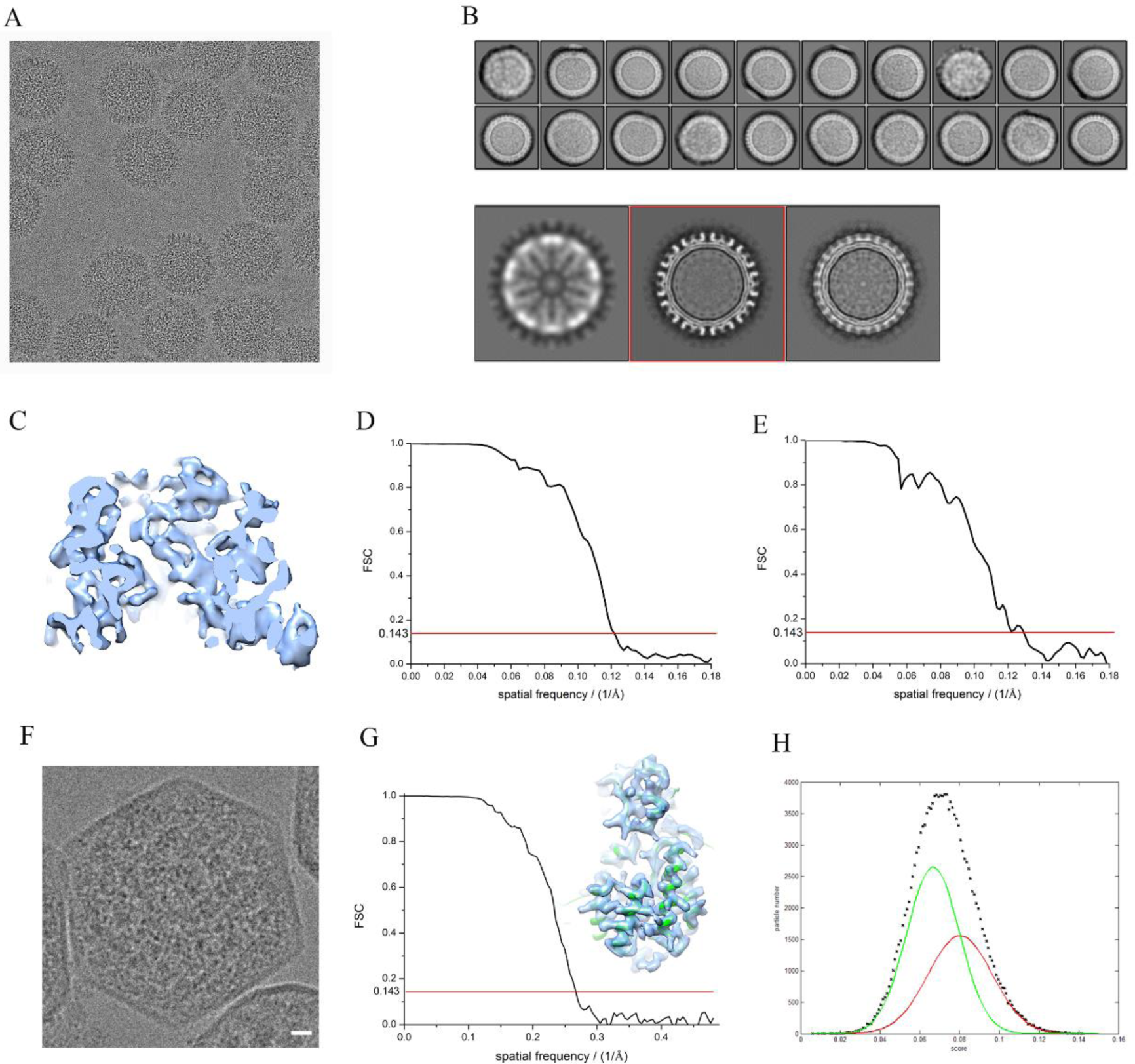
Application. (A) A micrograph of Bunyavirus particles. (B) Upper panel: 2D classification of viral particles binned by 4. Lower panel: 3D classification of viral particles selected from 2D classification. (C) Densities in the 7.7 Å map of pentamer. (D) FSC curve shows the resolution of hexamer map. (E) FSC curve shows the resolution of pentamer map. (F) An image of a carboxysome, Rubiscos are packaged inside; scale bar represents 10 nm. (G) The 3.7 Å map of Rubisco reconstructed using our isSPA method (right) and the FSC curve showing the resolution of the 3.7 Å map at gold standard (left). (H) Distribution of particle numbers to scores fitted by two Gaussian functions.

In the second sample, carboxysomes were purified from *Halothiobacillus neapolitanus* and the cryo-EM data were collected on this sample. The size of the carboxysomes ranged from 100 nm to 150 nm. We also purified Rubisco from fractured carboxysomes by adding extra freeze-thaw cycles before the centrifugation step for carboxysome purification. We collected the cryo-EM data of purified Rubisco, and obtained a 2.7 Å map of this complex (around 500 kD). This complex was then used as a model to pick the Rubisco inside the carboxysome in the cryo-EM images. The frequencies between 1/100 Å^-1^ to 1/8 Å^-1^ were used for picking. From the collected micrographs, ∼5,200 carboxysome images were extracted, and each image was cross-correlated with the 2.7 Å model at a frequencies range from 1/100 Å^-1^ to 1/8 Å^-1^. This procedure yielded a large group of locations and orientations, from which on average 150 solutions were picked per carboxysome for further processing. The non-alignment 3D classification was performed using RELION. Approximately 150,000 particles were selected, and a further auto local refinement with angular sampling of 0.9 degrees and translational sampling of 1.04 Å in RELION reported a 4.3 Å resolution map. To remove the false positive detections, the refined particles were sorted with phase residual using frequencies from 1/8 Å^-1^ to 1/5 Å^-1^. We tested three cutting thresholds (0.07, 0.08 and 0.09) and selected ∼82,000, ∼46,000 and ∼21,000 particles to the second-round refinement individually, resulting in 4.0 Å, 3.9 Å and 3.9 Å resolution. Further CTF refinement subsequently improved the resolutions to 3.9 Å, 3.9 Å and 3.7 Å. According to the fitting of two Gaussian distributions (Fig. 5H), the portion of true solutions at threshold 0.09 was estimated to ∼90%.

The success on Rubisco data set may owe to the high symmetry (D4) and its high abundance. We have shown above that our work flow can solve protein picked with precision below 0.1. According to the description of section 3.1, the recall of the 960kD protein with 200 times decreasing of abundance keeps the similar as a 480kD protein at precision 0.1. Thus, it is possible that a ∼1MD protein in Carboxysome thickness (∼120 nm) with 200 times lower abundance than that of Rubisco can be solved at a pseudo atomic resolution.

## 3. Discussion

In this work, we showed that the image-processing routines of isSPA method can resolve proteins (larger than 400 kD) at high resolution *in situ* with non-cryo-sectioning data when the thickness of the sample is around 120nm. The ability of finding the initial rotational and translational parameters of the target protein is closely associated with the size of the target protein and the thickness of the cryo-EM sample. Our recent results show that 50% of ∼350kD membrane proteins can be found and reconstructed to pseudo atomic resolution on 50nm-diamter liposomes. We are also able to reconstruct a 1.2MD membrane protein containing only 150kD soluble domain staying on the original cellular membrane formed liposome with an average diameter of ∼150nm at high resolution. Thus, isSPA method can very efficiently help to solve structures of different kinds of membrane proteins on liposome constituted by their native lipid membrane. Consdering the thickness of cryo-sectioning sample and the abundance of the target protein, it is worth to try on a protein with size ∼ 1MD using this method. Low SNR is the major obstacle of determining the initial parameters of protein complexes. Thus, this method will benefit greatly from hardware improvement such as better direct detectors or new generation of phase plate that improves the SNR of images at frequencies ranging from 1/20 Å^-1^ to 1/8 Å^-1^ (0.1∼0.25 Nyquist at a pixel size of 1 Å) or better tool to obtain a thinner cryo-sectioning.

When the protein is imaged in the in situ environment, the overlapping densities ruin the SNR of low frequency signals of the target protein in the images. The SPA algorithms improve the 3D structure iteratively from a low resolution 3D map to a high resolution 3D map. However, such a method become invalid when the low frequency signals of the target protein was ruined by overlapping densities. Thus, routine none-reference 2D classification and 3D classification starting from a low resolution initial model usually converge to wrong results. Thus, starting from the local refinement in both structure refinement and 3D classification is strongly suggested in isSPA. The structural information of a template with frequency above 1/8 Å^-1^ is involved in picking the particle, therefore the FSC curves below 1/8 Å^-1^ is not calculated from two completely independent datasets. However, the FSC beyond 1/8 Å^-1^ is not affected by the template and can be considered as gold-standard.

Generally, isSPA is not a good choice for mapping protein complexes in in situ environment due to the relatively low detection efficiency. Moreover, the z height of the protein in the in situ environment is determined by its defocus value in the micrograph. Although, the per-particle CTF refinement improves the defocus value of the protein, error remains. It prevents an accurate localization of the target protein along Z direction. Thus, if the distribution of target protein is unknown, we suggest that a tomographic study prior to isSPA can be used to show the distribution of target protein in the in situ environment and provide a medium resolution template if it is necessary. Then, isSPA can be used to further improve the resolution in an efficient way.

## 4. Materials and Methods

### 4.1. Reducing the false positive results by sorting

The introduction of high resolution (1/8 Å^-1^) information in global detection will generate a large amount of false positive detections due to the especially low SNR. These false positives are presented as noise beyond 1/8 Å^-1^ and produces biased features in the reconstruction below 1/8 Å^-1^ which affect the alignment against the reconstruction, thus prevent from pushing high resolution in the following refinement procedures. To further reduce the false positive results, we sorted the selected images based on a different score. The signals with frequency range beyond 1/8 Å^-1^ are recommended for calculating the score since they are less affected by biased feature.

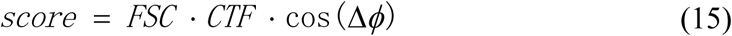

Due to the extremely low SNR in high frequencies range, the constant n is much smaller than *1/SSNR*, thus we ignored the overlapping density and used *FSC* and *CTF* as weighting factors.

### 4.2 Workflow of isSPA method

IsSPA is a single-particle like method to reconstruct *in situ* structures from untilted images. The images therefore contain overlapping protein densities, and the workflow includes: (1) estimate SSNR parameters by fitting power spectrum with two exponential functions (one for noise and the other for signal); (2) for particle picking, raw images are filtered by our optimized weighting function with known SSNR parameters, then the filtered images and projections of 3D model are input to program; (3) the output of program presents a large group of locations and orientations, thus we merge the neighbor solutions according to their translations and rotations; (4) we sort solutions from each image by cc value, and the number of solutions are selected depending on the estimated particle number in each image; (5) scripts were used to extract potential particles and perform a local refine and 3D classification on these particles using different cryo-EM software packages such as RELION, JSPR and so on; (7) a sorting algorithm was provided to remove false positive particles; (8) the remaining particles are performed local refinement to achieve the final reconstruction.

### 4.3 Image acquisition of test data

Images of Herpes simplex virus type 2 particles (HSV-2) (∼120 nm in diameter) and Reovirus particles (∼90 nm in diameter) obtained from previous studies were analyzed for testing. HSV-2 data were collected on an FEI Titan Krios microscope operated at 300 kV equipped with a Falcon2 camera, with pixel size of 1.38 Å/pixel and a total dose of ∼25 e^-^/Å^2^. Reovirus data was acquired on an FEI Titan Krios microscope operated at 200 kV with a Gatan K2 Summit camera, and the pixel size is 1.04 Å/pixel with a total dose of ∼30 e^-^/Å^2^.

### 4.4. Image acquisition of Bunyavirus

Bunyavirus data were collected on an FEI Titan Krios EM operated at 300 kV equipped with a Gatan K2 Summit detector. Data collection was performed with Serial EM using a nominal magnification of 22,500 with a total dose of ∼50 e^-^ resulting in a pixel size of 1.3 Å/pixel. Each micrograph was recorded as a movie composed of 36 frames. A total of 1,849 such movies were collected and 10,950 viral particles were extracted from motion-corrected micrographs.

### 4.5. Image acquisition of Carboxysome

Purified carboxysomes were applied to a glow-discharged Quantifoil R1.2/1.3 300-mesh copper holey carbon grid (Quantifoil, Micro Tools), blotted under 100% humidity at 4°C and plunged into liquid ethane using a Mark IV Vitrobot (FEI). Micrographs were acquired on a Titan Krios G2 microscope (FEI) operated at 300 kV with a K2 Summit direct electron detector (Gatan). A calibrated magnification of 130k times was used for imaging, yielding a pixel size of 1.04 Å on images. The defocus was set at 1.5 to 2.0 μm. Each micrograph was dose-fractionated to 32 frames under a dose rate of 8 e^-^/pixel/s with a total exposure time of 10.4 s.

## Acknowledgements

We thank L. Kong for cryo-EM data storage and backup and Z. Y. Lou in Tsinghua University for offering us the cryo-EM data of Bunyavirus. Cryo-EM data collection was carried out at the Center for Biological Imaging at the Institute of Biophysics (IBP), Chinese Academy of Sciences (CAS). We thank X. Huang, B. L. Zhu, G. Ji and other staff members at the Center for Biological Imaging (IBP, CAS) for their support in data collection. The project was funded by the National Key R&D Program of China (2017YFA0504700), the Natural Science Foundation of China (31930069) the Strategic Priority Research Program of the Chinese Academy of Sciences (XDB37040101) and the Key Research Program of Frontier Sciences at the Chinese Academy of Sciences (ZDBS-LY-SM003), X.Z. received scholarships from the ‘National Thousand (Young) Talents Program’ from the Office of Global Experts Recruitment in China.

## Abbreviations

SNR: signal to noise ratio
SSNR: spectrum signal to noise ratio
CTF: contrast transfer function
cc: cross correlation
FSC: Fourier Shell Correlation
HSV: Herpes simplex virus
SPA: single particle analysis

## Compliance with ethics guidelines

Jing Cheng, Bufan Li, Long Si and Xinzheng Zhang declare that they have no conflict of interest. This article does not contain any studies with human or animal subjects performed by the any of the authors.

## Supplementary Materials

**Figure S1.**
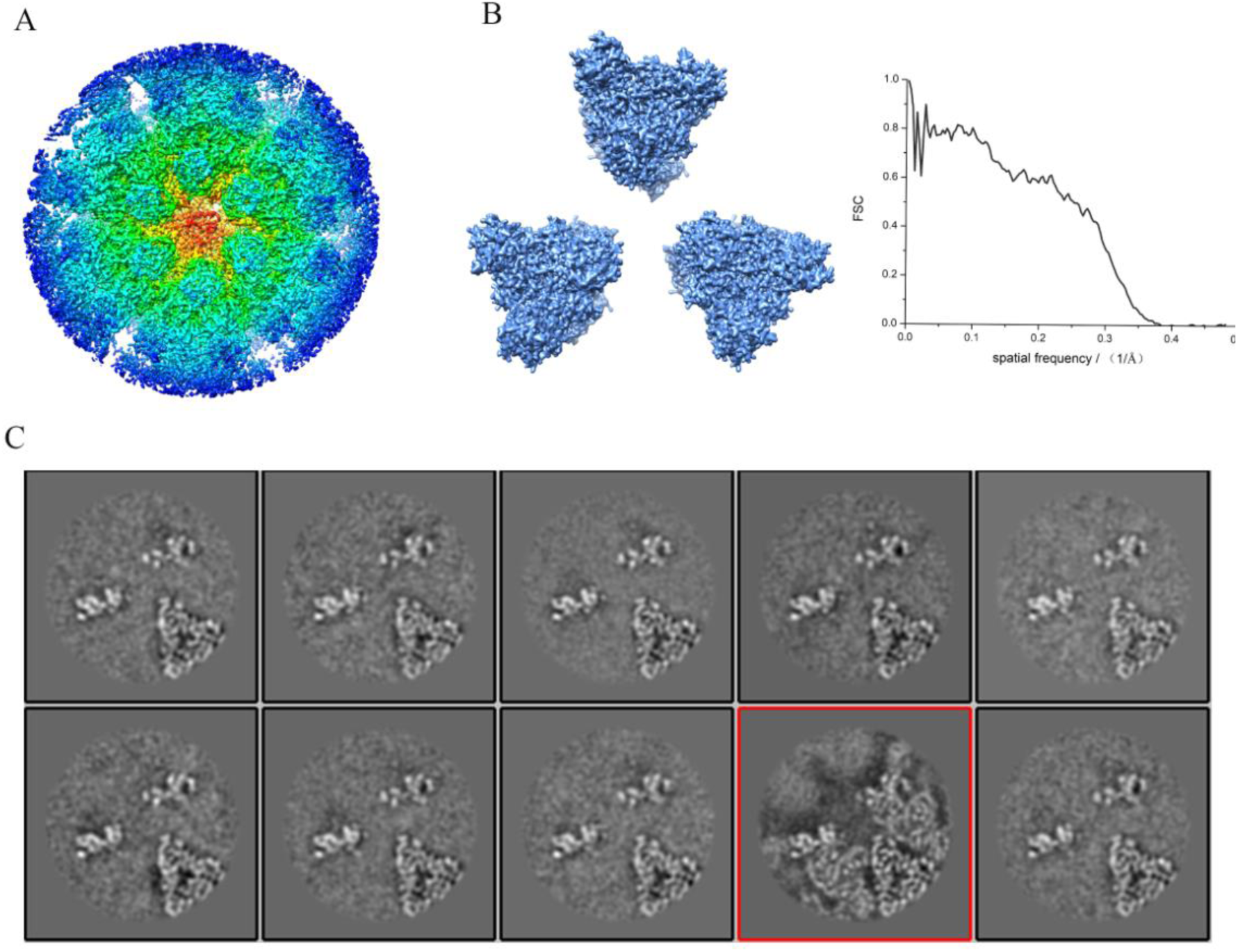
Data processing of Reovirus. (A) The 4.0 Å reconstruction of reovirus hexamer by our isSPA methods with previous reconstruction as picking model. (B) Left: EM map produced by pdb of the crystal structure (1jmu). Right: FSC curve describing the similarity between the target and homologous model. (C) 3D classification of selected particles from global search with homologous map as picking model, and particles in class 8 (which is bolded in a red square) were most true detections.

**Figure S2.**
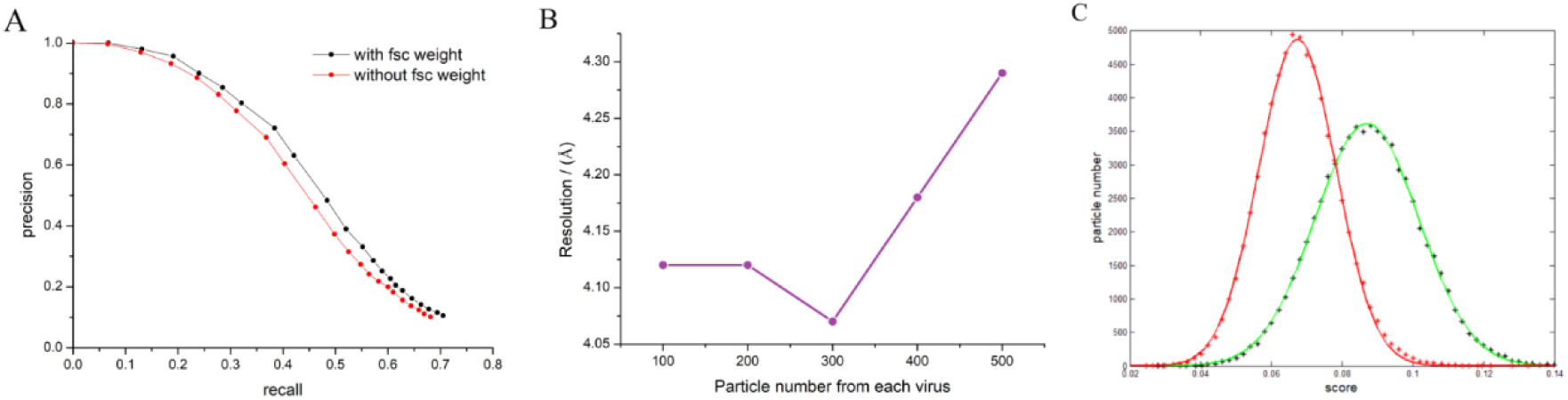
FSC weight and picking threshold on HSV-2 dataset. (A) Precision-recall curves of the ∼2 MD protein rescaled to ∼8 Å at FSC 0.5. We tested on HSV-2 with FSC weighting (black) and without FSC weighting (red). (B) The relationship of resolution to threshold tested on HSV-2 dataset before PR sorting (black) and after PR sorting (purple). (C) Gaussian fitting of particle number to score distributions by true solutions (red) and false solutions (green).

**Figure S3.**
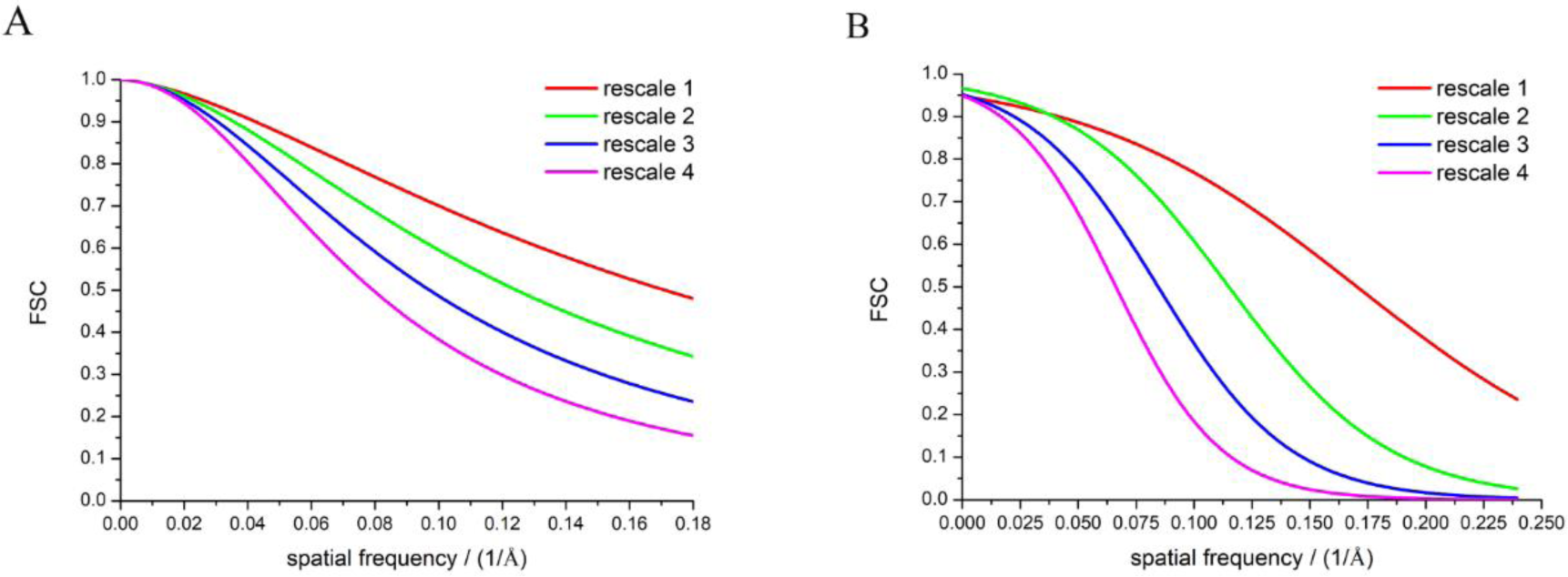
FSC curves of rescaled maps with the origin maps. (A) HSV-2 hexamer rescaled at different scales. (B) Reovirus hexamer rescaled at different scales.

**Figure S4.**
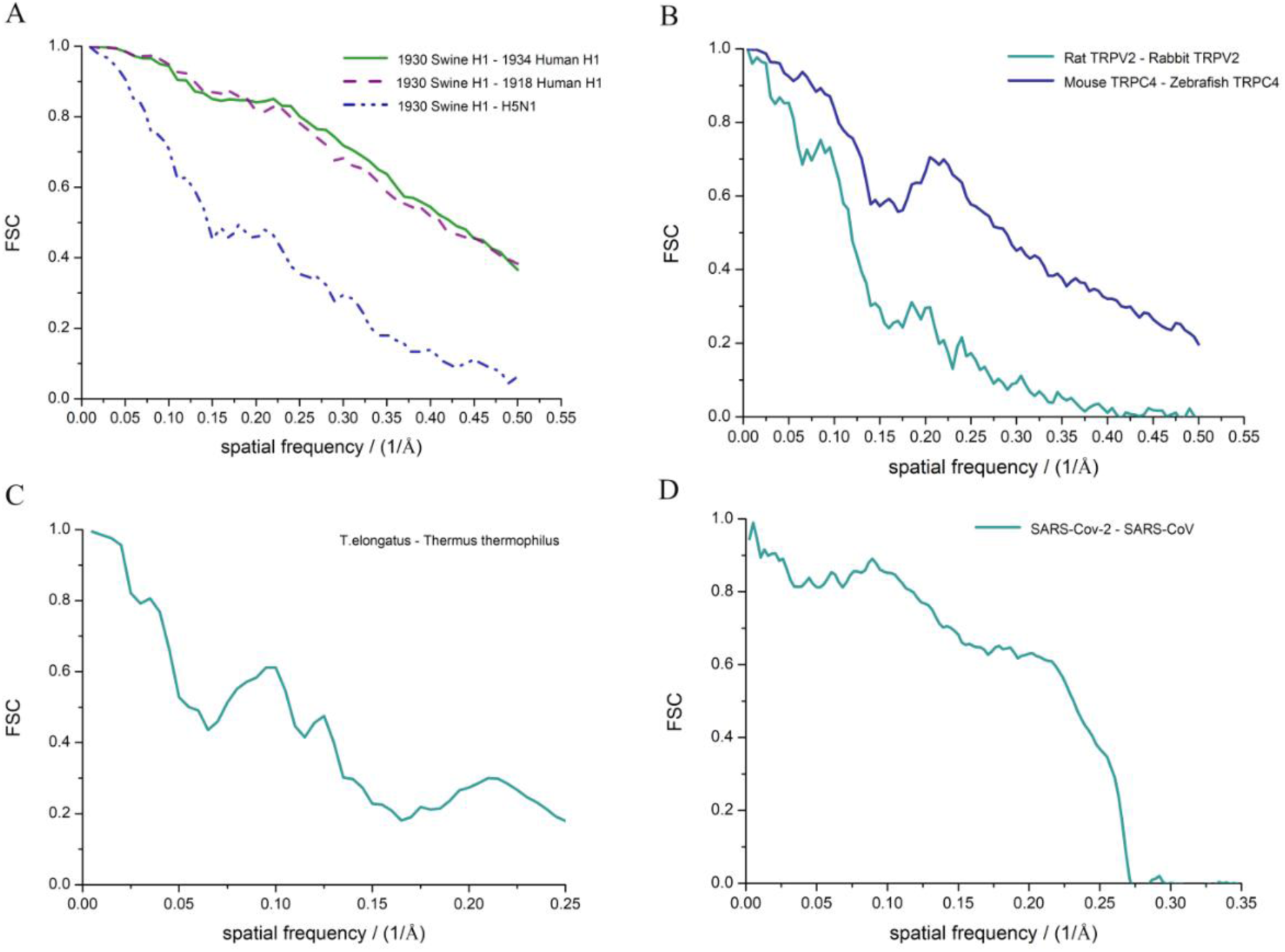
FSC curves between homologous structures. (A) Glycoproteins of influenza viruses. (B) Ion channels proteins. (C) NDH. (D) Spike proteins of coronaviruses.

**Figure S5.**
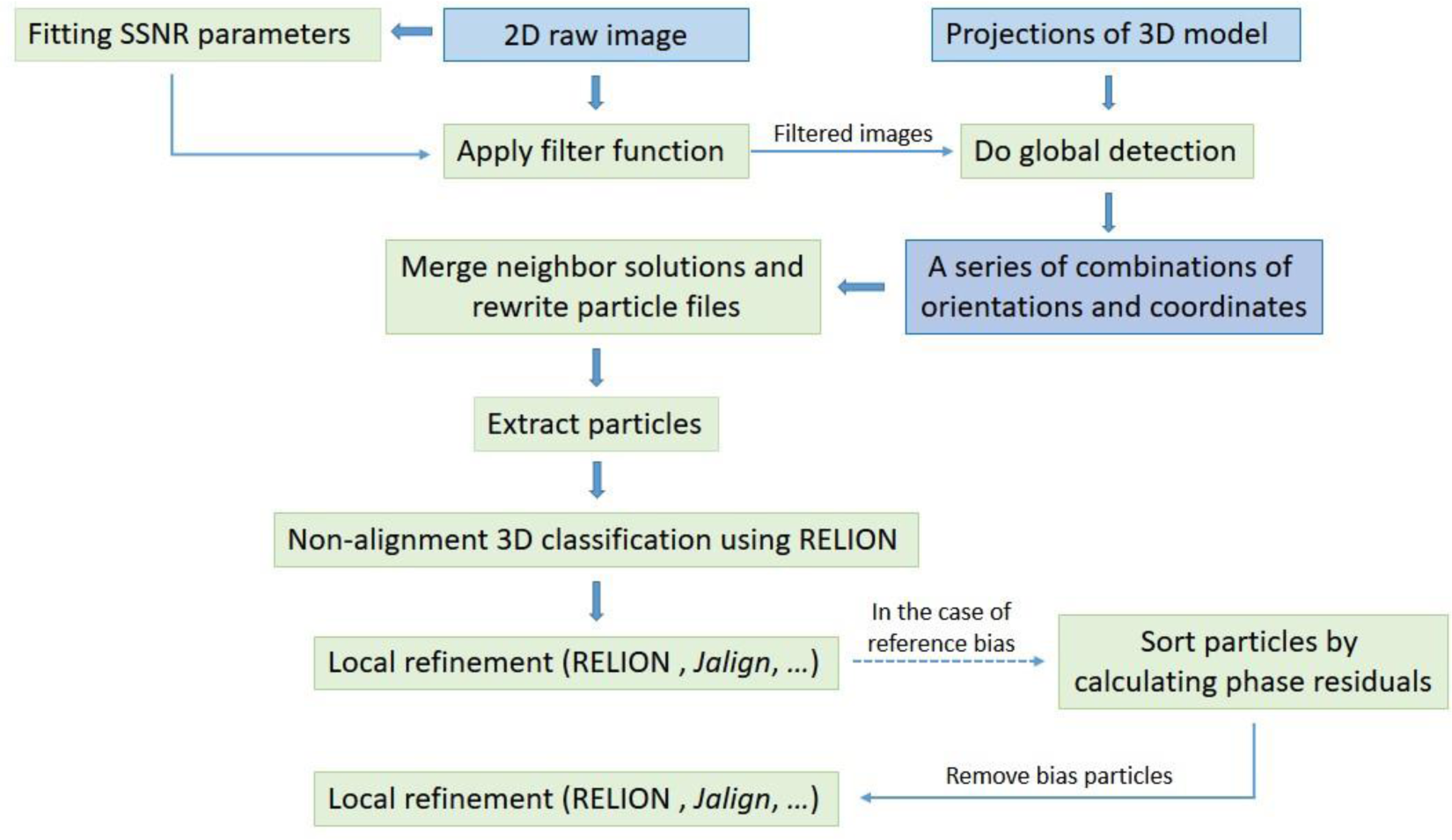
Flowchart of isSPA method.

**Figure S6.**
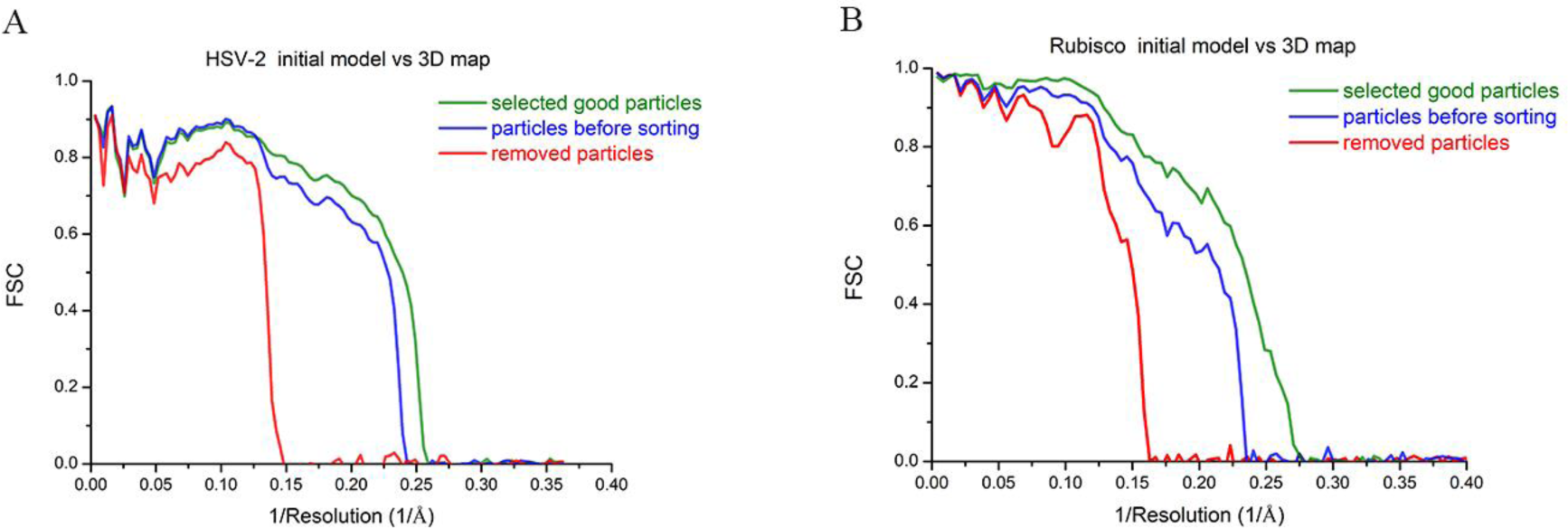
FSC with high resolution initial model. (A) FSCs of 3D reconstructions by selected particles according to sorting (green), removed bad particles according to sorting (red) and both particles before sorting (navy) with high resolution initial model. (B) the same processing with Rubisco images as in (A).

